# Batch-Corrected Distance Mitigates Temporal and Spatial Variability for Clustering and Visualization of Single-Cell Gene Expression Data

**DOI:** 10.1101/2020.10.08.332080

**Authors:** Shaoheng Liang, Jinzhuang Dou, Ramiz Iqbal, Ken Chen

## Abstract

Clustering and visualization are essential parts of single-cell gene expression data analysis. The Euclidean distance used in most distance-based methods is not optimal. Batch effect, i.e., the variability among samples gathered from different times, tissues, and patients, introduces large between-group distance and obscures the true identities of cells. To solve this problem, we introduce Batch-Corrected Distance (BCD), a metric using temporal/spatial locality of the batch effect to control for such factors. We validate BCD on a simulated data as well as applied it to a mouse retina development dataset and a lung dataset. We also found the utility of our approach in understanding the progression of the Coronavirus Disease 2019 (COVID-19). BCD achieves more accurate clusters and better visualizations than state-of-the-art batch correction methods on longitudinal datasets. BCD can be directly integrated with most clustering and visualization methods to enable more scientific findings.

## 1 Introduction

Gene expression reflects the identity of a cell. Single-cell RNA sequencing (scRNA-seq) technologies profile thousands of cells simultaneously [1], enabling trajectory inference to reveal the course of cell development and transformation [2]. Although large amount of data have been gathered from different tissues among large cohorts of patients [3], technical variance among separately assayed samples often overshadow the similarity of cells, resulting in a less meaningful disconnected trajectories (Figure 1a). In addition, nuisance differences among individual participants also blur the shared underlying biology. These phenomena, often referred to as batch effects, complicate the analysis of single-cell data. For samples collected from the same condition (i.e., same time and tissue but different participants), all the differences among samples may be deemed as batch effect and get corrected. Nevertheless, over-correction is reported [4], any can be a greater problem on longitudinal data, where true biological changes and nuisance factors are entwined (Figure 1c).

**Figure 1:**
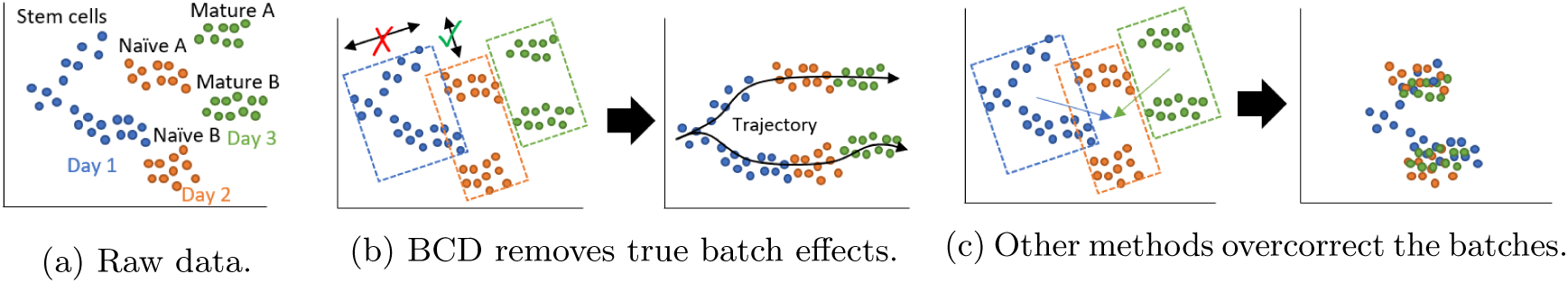
Illustration of how BCD handles batch effect differently.

As a practical example, researchers have collected data from the developing retina of 13 mice at different ages [5]. Ideally, the embedding of such data should reveal a fluent trajectory of how pluripotent stem cells differentiate/evolve into multipotent stem cells, and finally to specific cell types. However, a mouse may not survive the tissue extraction, and thus each sample in the dataset is from a unique mouse. Batch effect exists among samples collected from different mice. In addition, there is no bijective (i.e., injective and surjective) mapping for cells from different samples. Thus, existing methods addressing batch effect in longitudinal data assume that the features are measured on the same set of entities (cells) at different time points [6] are not applicable in this scenario. Besides, because single-cell profiling technologies consume a cell when measuring it, the cells would not be the same or matched even if the samples were collected from the same tissue from the same mouse.

To date, multiple methods have been published to correct batch effects [7]. For example, Seurat utilizes mutual nearest neighbor (MNN) to identify similar clusters in different batches and integrates those batches by removing the differences [8]. Harmony also integrates proximal clusters, but in an iterative way through soft clustering [9]. Liger uses nonnegative matrix factorization (NMF) to separate common and sample-specific features [10]. A neural network approach, scVI, combines variational autoencoder and zero-inflated model to visualize and correct the data. Notably, Harmony generates integrated clusters and visualization without giving a corrected expression profile. It is not deemed a significant drawback, however, because corrected datasets are often hard to interpret. Researchers usually only cluster and visualize the data using such methods, and recur to statistical tests that control for the batch effect in the downstream analyses [11]. These methods have been utilized in a few large-cohort studies and satisfyingly removed the batch effect among samples [8, 9]. However, none of them are designed for longitudinal data.

In an effort to address this issue, we noticed the “locality” among the single-cell samples. Specifically, samples gathered in proximal time points are expected to be more alike than from distant times. Similarly, the later two cell types are differentiated in development, the more alike they will be. That means the difference between two samples is more likely to be nuisance if those samples are collected from the same time point or adjacent time points. Here, we define a new distance function, termed Batch-Corrected Distance (BCD) which exploits the locality to precisely remove the batch effect (Figure 1b) but keep biologically meaningful information that forms the trajectory. We show the derivation of BCD in section 2. As a distance metric, it is naturally compatible with all state-of-the-art distance-based clustering and embedding methods. We compare BCD with Euclidean distance, Seurat integration, and Harmony on simulated data and a mouse retina development data, and show applications of BCD on COVID-19 patient data and human fetal lung development data in Section 3. The results clearly show the benefit of BCD to biology studies.

## 2 Methods

### 2.1 The Batch-Corrected Distance

Trajectory inference, in general, aims to find a graph *G* = (*V, E*) which reflects the hop-by-hop gradual change (*E*) of cells (*V*_*i*_, *V*_*j*_, … ∈ *V*) by optimizing

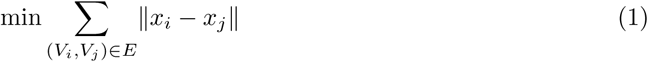

subjecting to a set of constraints [2, 12]. Here, *x*_*i*_ is the profile of cell *i*, which can be the whole gene expression, or the first few principal components (PCs). Eulidean distance is the most widely used metric, while the Mahalanobis distance

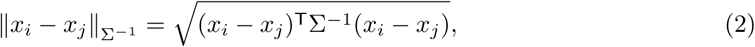

where Σ is the covariance matrix calculated from all the samples, may be used to account for different (co)varainces among features. To account for the batch effect, we propose to redefine the Σ as

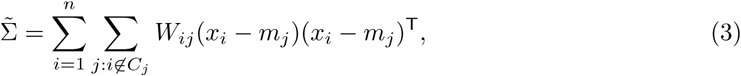

where *C*_*j*_ = {*i* |cell *i* is in batch *j*} is the batch information and *m*_*j*_ is the centroid of each batch. If weight *W*_*ij*_ ≡ 1, it degrades to the metric defined by Qi and Davidson [13] for generating an alternative clustering different from the original *C*. In essence, it removes the variances across the batches, while retain the variance within each batch. For a longitudinal dataset, each sample is (and all the cells in the sample are) collected from a time point, denoted as *t*_*j*_ (and *τ*_*i*_). To utilize the temporal/longitudinal locality, we set *W*_*ij*_ to be

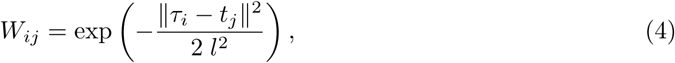

where *l* (set to 1 in our experiments) is the length scale within which two samples are considered temporally/spatially close. The covariance of proximal time points are weighted more, and thus are suppressed in the refined distance. When the dataset contains both temporal and spatial labels, *τ*_*i*_ and *t*_*j*_ can be vectors that include both labels. Inversed Cholesky-decomposed 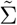 can be used to transform the data. If first k PCs are used, the computational complexity is *O*(*nk*^2^ + *k*^3^).

### 2.2 Gene expression data processing

We use Seurat [14], an R package, to analyze the gene expression data. The package provides functionalities to normalize data, find highly variable features (i.e., genes) by variance stabilizing transformation, scale the features, perform principal component analysis (PCA), and visualize the result with UMAP (uniform manifold approximation and projection). This is the de facto standard single-cell data analysis protocol. The normalization step, in specific, normalizes the summation of gene expression in each cell to be one. The scaling step standardizes each gene so that the average expression over all cells is zero, and the standard deviation is one. UMAP is a nonlinear embedding method visualize data by their distance [15].

Also provided in Seurat is a data integration method that corrects batch effect [8]. It first projects samples into a common subspace using canonical correlation analysis (CCA), and then finds MNNs in the CCA subspace as “anchors” to correct the data. We refer to it as Seurat integration (not to be confused with the entire Seurat protocol). Harmony first projects the data into a lower-dimensional PCA space, and then iteratively removes batch effects. At each iteration, it clusters cells while maximizing the diversity of batches within each cluster and calculates a correction factor for each cell to remove the batch effect. Tran et al. [7] show by systematic assessments that both methods are state-of-the-art. Thus, we compare BCD with them. To ensure good comparability, we implemented BCD with an interface to Seurat. This choice also makes BCD easy to use for biology researchers familiar with Seurat.

## 3 Results

### 3.1 Simulation study

To validate BCD, we simulated a dataset with seven samples. We assume that all cells are differentiated from an initial state, stem cells named S. S differentiates to two multipotent cell types A and B, which differentiate to terminal types A1, A2 and B1, B2, respectively (Figure 2a). We simulated 200 genes, and assigned a unique ideal gene expression profile for each cell type, denoted as *s, a, b, a*_1_, *a*_2_, *b*_1_ and *b*_2_. Because the development of the cell types is gradual [16], we used weighted mean to represent the process in the simulated samples. For example, to simulate intermediate states between A and A1, we used (*θa* + (1 − *θ*)*a*_1_) as its profile, where *θ* is set to be uniformly distributed in a range corresponds to the developing stage of a sample (Table 1). The ambiguous cell types (*θ* close to 0.5) are denoted as “A->A1”, while cells more similar to “A” or “A1” (*θ* close to 0 or 1) are labeled as them each.

**Table 1:**
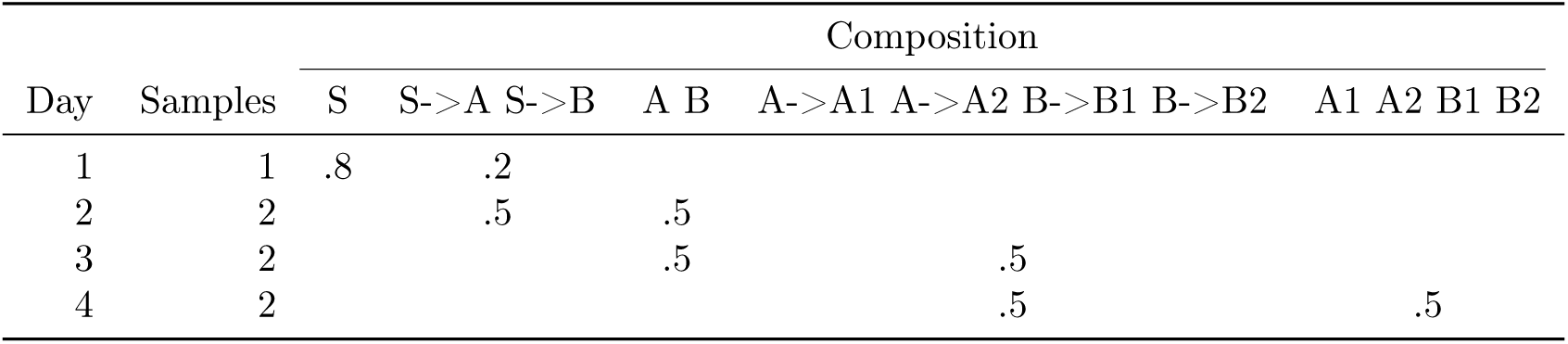
Simulated dataset

**Figure 2:**
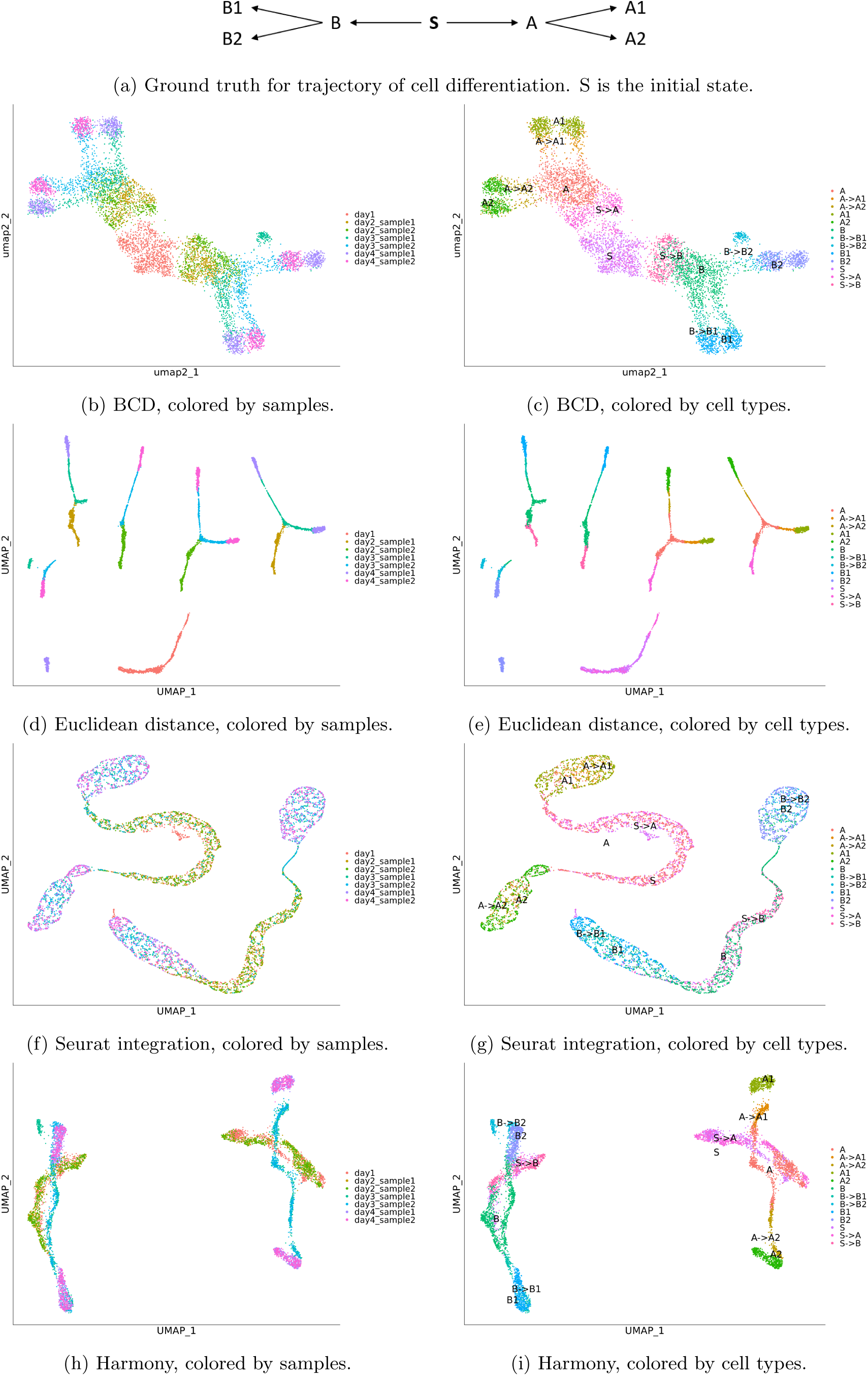
Results for the simulated data. Left panels are colored by samples. Right panels are colored by cell types. Each row represents a method.

We randomly selected a set of genes to be “susceptible to the batch effect”, and added different additive components on those genes in different samples. For every single cell, we further added random noise into the ideal profile and used a Poisson distribution to generate the gene expression. The number of samples at each time point, and compositions of samples are shown in Table 1. It can be seen that the cell types gradually evolve over time.

We used Seurat to process the data with BCD, Euclidean distance, Seurat integration, and Harmony. In the process, 150 highly variable genes are selected, and 30 PCs are used. The result of BCD are shown in Figure 2b and 2c. Given the prior knowledge that all cell types originate from S, it can be clearly seen that it branches to A and B, and further to A1, A2, and B1, B2, respectively. Small batch effect can still be seen within A1, A2, B1, and B2, but are largely mitigated. The samples are also well organized by their time. Day 1 is at the center, day 4 is separated to A1 and A2 at the top-left and B1 and B2 at the bottom-right corner, while day 2-3 corresponds to the trajectory of which the cells differentiate.

In contrast, Euclidean distance, the baseline, does not correct for any batch effect (Figure 2d and 2e). Consequently, the terminal types A1, A2, B1, and B2 each splits into two groups, corresponds to different samples. It also fails to illustrate the evolution of the cell types. Seurat integration shows high capacity of removing batch effects, as the cells from multiple days are mixed together (Figure 2f). However, it is an over-correction. For example, S, S->A and A are mixed together, and so do A->A1 and A1, and A->A2 and A2. The same applies to the branch of B. It also leads to incorrect trajectory inferences, as A is now closer to A than A->A1. Overall, it makes it harder to delineate the cell differentiation (Figure 2g). The result of Harmony is slightly better (Figure 2h and 2c). The two batches are fairly mixed for A1, A2, B1, and B2. The trajectory of S->A to A, then to A->A1 and A->A2, and finally to A1 and A2 can be seen, although the B->B2 and B2 are misplaced. Besides, the branch of A and branch of B are still in disconnected clusters, leaving their relationship with S undiscovered. Overall, BCD performs the best in the four methods. For time consumption, BCD uses less than 1 second, while Harmony uses 9 seconds and Seurat integration uses 20 seconds.

### 3.2 Mouse retina development dataset

We applied BCD to a dataset including 13 single-cell specimens collected from developing retina of one sample at E12 (day 12 embryo), two at E14, one at E16, two at E18, one at P0 (postnatal day 0), two at P2, one at P5, two at P8, and one at P14 [5]. A total of 110,359 cells are collected and process with Seurat, where 2,000 highly variable genes are selected, and 30 PCs are used.

The development of mouse retina is well-understood by the field. Briefly, retinal progenitor cells (RPCs, including early RPCs and late RPCs) differentiate into Neurogenic cells, which further differentiate into photoreceptor precursors, Amacrine cells, horizontal cells, and retinal ganglion cells [17]. The photoreceptor precursors differentiate into cone cells, rod cells, and bipolar cells [18]. These cell types are all neural cells. Late RPCs also differentiate to Müller glia [17]. These cells form a neural network where each terminal cell type has a unique function. Cone cells and rod cells forming the input layer are photoreceptors that work in light and dark environment, respectively. The signal from cone cells and rod cells propagate through the bipolar cells first and then retinal ganglion cells to go to the brain. Horizontal cells provides horizontal connections between photoreceptors and bipolar cells, while amacrine cells perform similar function between bipolar cells and retinal ganglion cells. These cells form two hidden layers. Müller glia is an auxiliary cell type that supports the aforementioned neural cells.

The result of BCD is shown in Figure 3b and 3c. There is a clear trajectory of the cells gradually evolving from E11 to P14. The aforementioned evolution trajectory of the cell types can also be seen along the trajectory. The Euclidean distance also performs relatively well on this dataset. However, clear batch effect can be seen in Figure 3d, where cells are clustered by samples. As a result, a cell type is divided into multiple clusters in Figure 3d, making it harder to delineate the differentiation trajectory of the cell types. Both Seurat integration and Harmony over-corrects the batch effect. Although the samples are well mixed (Figure 3f and 3h), many cell types are mixed. For example, early RPC, late RPC, and Müller glia are not distinguishable from the result, and so do horizontal cells, amacrine cells, bipolar cells, and retinal ganglion cells (Figure 3g and 3i). Even worse, the bipolar cells, which should be differentiated from photoreceptor precursors, are misplaced to directly under neurogenic cells. These mistakes are because these methods do not utilize the temporal locality of the samples. For this dataset, BCD uses less than 2 seconds, while Harmony uses 5 minutes. Seurat integration costs more than 3 hours.

**Figure 3:**
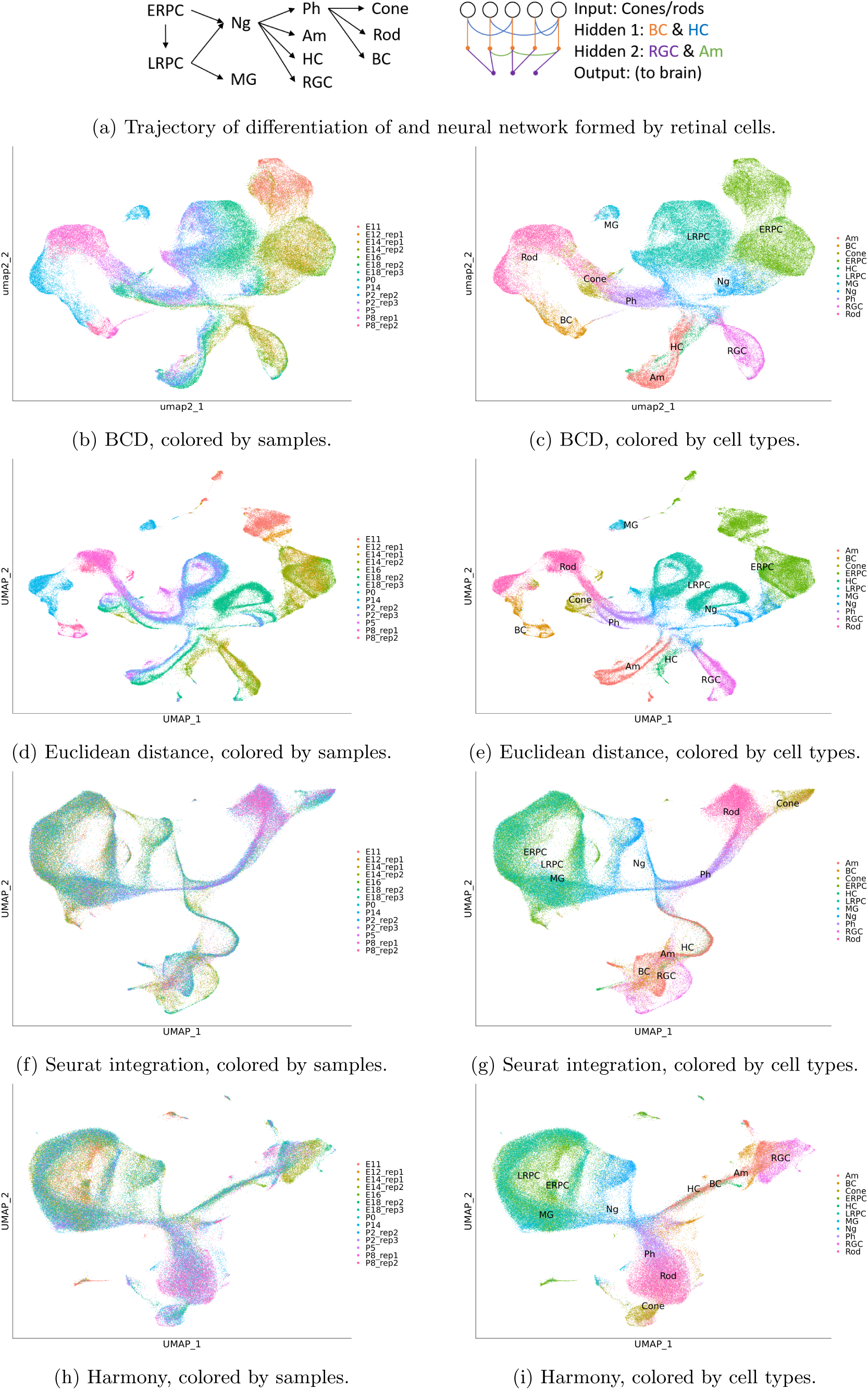
Results for the retina development dataset. ERPC: early retinal progenitor cell, ERPC: late retinal progenitor cell, MG: Müller glia, NG: neurogenic cell, Ph: photoreceptor precursors, Am: Amacrine cell, HC: horizontal cells, RGC: retinal ganglion cells.

### 3.3 COVID-19 immune compartment datasets

The global pandemic COVID-19 has reportedly infected 6.3 million people worldwide, with the death toll at 380 thousand. Understanding the immune response to the virus is essential to developing treatments. We applied BCD to a recently published dataset including immune compartment samples collected from 13 participants [19] including 4 health controls (HC1-4), 3 moderate (M1-3) cases, and 6 severe (S1-6) cases. Although these are all different participants, we consider HC, M, and S a time course to reflect the progression of the disease. We used Seurat and BCD to process the data. During the process, 2,000 highly variable genes are selected, and 30 PCs are used.

The result is shown in figure 4. The largest group is macrophages, which recognize and destroy virus-infected cells. The result shows from top to bottom a gradual change from those in health controls, to moderate cases, and to severe cases. In contrast, Euclidean distance yields disconnected groups of macrophages because of the batch effect, while Seurat integration and Harmony overcorrect the effect and confuse cells from moderate cases with those from health controls (Supplementary Material). The changes in gene expression that drive the differences can be attained by differential expression analysis. Similar trajectories can also be seen on T cells and plasma cells, which kill infected cells and produce antibodies, respectively. Using these pieces of information, researchers can identify the most effective form of immune cells and find ways to transform others into it to treat the disease.

**Figure 4:**
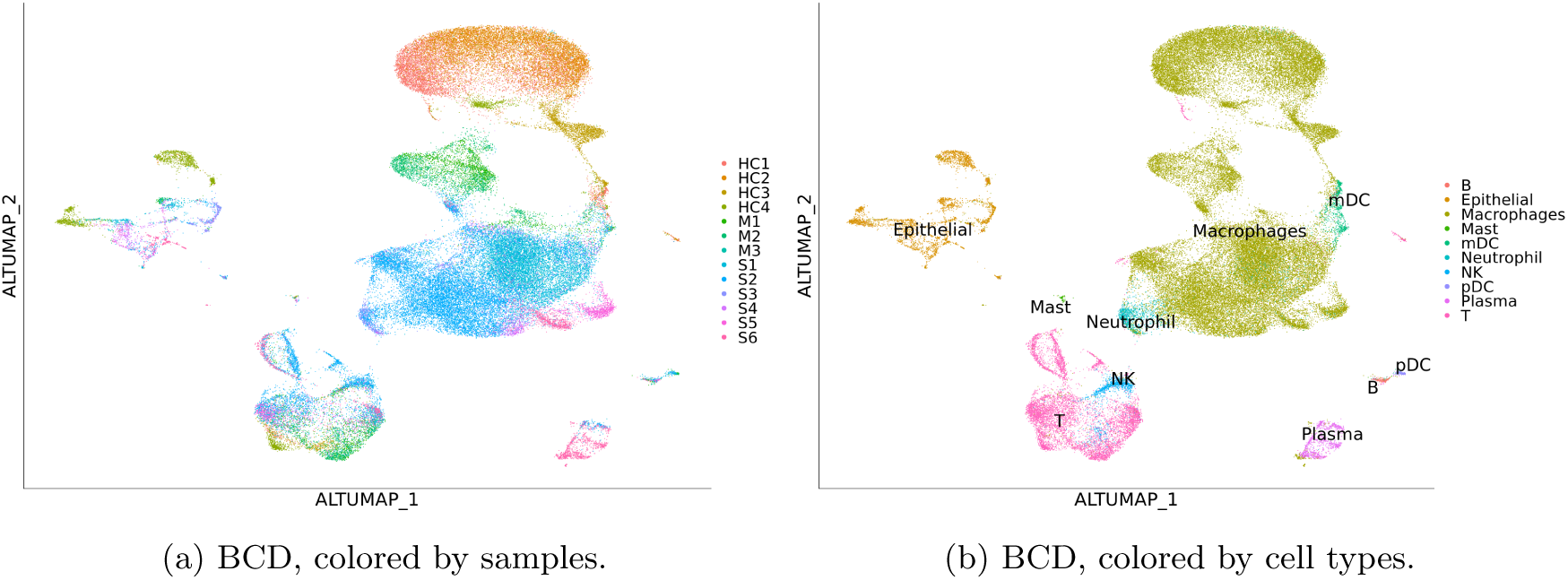
Results for the COVID-19 dataset. Results of other methods are in Supplementary Material.

### 3.4 Human Lung datasets

Besides studies of the immune compartment, knowledge of lung development may also help cure the disease and restore the functionality of the lung [20]. Miller et al. has produced a single-cell dataset of cells from [21]. Cells from fetal human lungs are collected at week 11.5 (W11.5), W15, W18, and W21, and are available for trachea, small airways in the lung, and the distal tip of the lung.

We explored the dataset using BCD. Because both temporal and spatial information of the samples is available, we use a vector [*week, location*] to label each sample, where *week* is the number of weeks mentioned above, and *location* is set to be 0, 2, and 4 for trachea, small airways, and distal lung, respectively. The result is shown in Figure 5. A branching trajectory can be seen from W11.5 to W18 for distal lung and small airways mesenchymal cells. The cells from the two locations are similar at the early stage (W11.5), but become more distinct when they are more developed (W15 and W18). In contrast, affected by the batch effect, Euclidean distance and Harmony show W15 and W18 small airway mesenchymal cells as isolated clusters, with no connection with W11.5, while Seurat mixes all mesenchymal cells across the two locations and all times points, blurring the trajectory (Supplementary Material). Similar trajectories also show for Epithelial cells, endothelial cells, and pericytes. Researchers may use these pieces of information to further study the changes in gene expression in the cell type developments and develop treatments.

**Figure 5:**
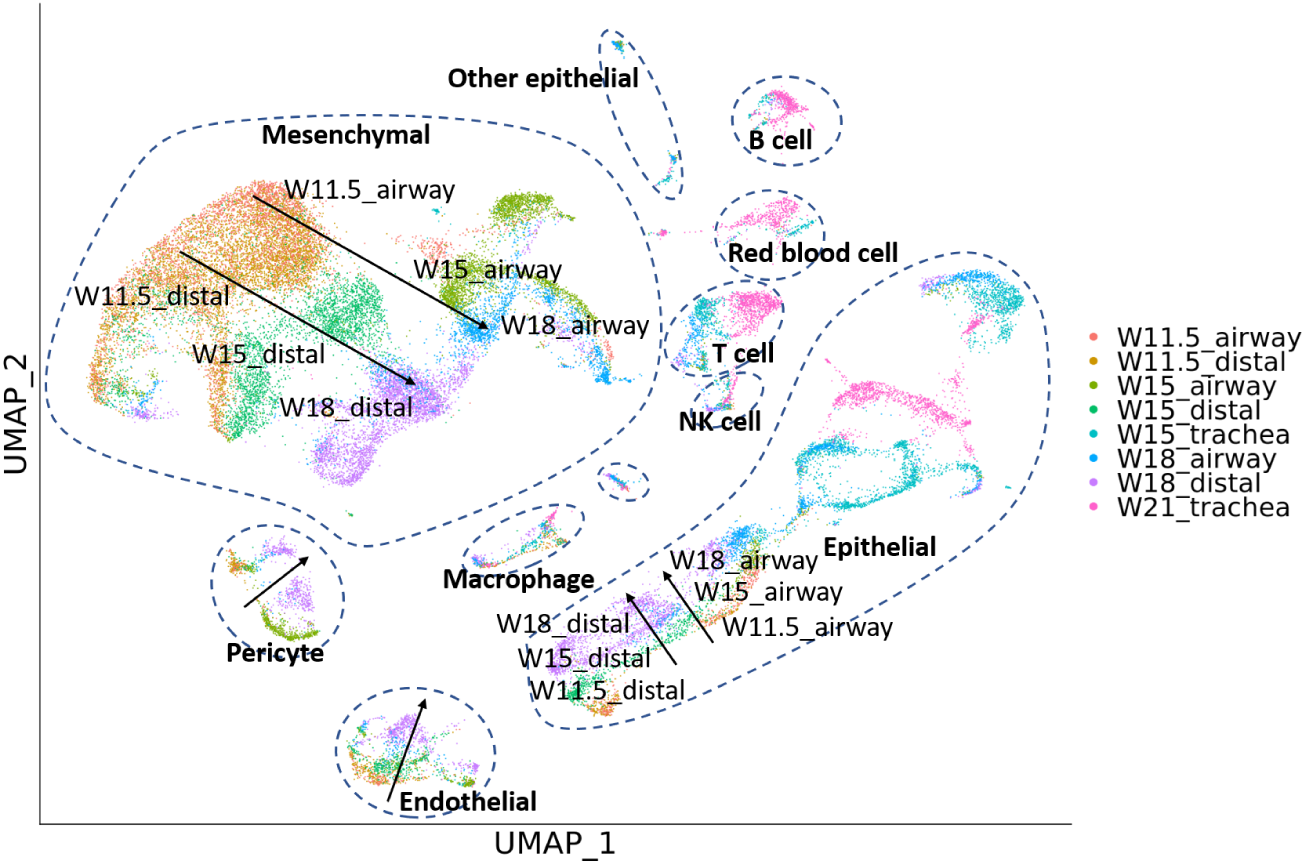
Results for the human fetal lung development dataset. Arrows are added for visual reference.

## 4 Discussion

In order to build a reliable trajectory of cell type development from a longitudinal dataset, the batch effect should be corrected for. This is a new problem as state-of-the-art batch effect correction methods cannot utilize the time/spacial information, and thus result in over-correction. Our method, BCD, utilizes such information to accurately identify and remove the batch effect, while preserving the correct structure of the data. The results based on the BCD clearly show the evolution of the cell types through time. The discovered trajectory helps researchers make hypotheses of the physiological changes during development or pathological changes as a result of disease infection. The changes can be ascertained by differential gene expression analysis.

BCD assumes that the batch effect is shared by all the samples. This is a reasonable assumption because many kinds of batch effects have a biological basis. For example, Cellular stress response stimulated by the sample preparation affects certain genes. In the case the set of genes do change, the local linear approximation may be used. Because the sums and products of distances are guaranteed to be valid distances, multiple BCDs each correcting for the batch effect in a specific set of samples can be combined as a consensus distance.

BCD is intrinsically a linear transformation, which may be insufficient for more complex batch effects including interactions of genes. Nevertheless, it is a proof of concept that the time/spatial locality should be considered in batch correction for longitudinal datasets, which are on the rise. Nonlinear methods may be invented based on the same concept. For example, kernelization can be a direct extension to BCD. Besides, BCD is very fast to run and can be used in the exploration of the dataset with negligible cost.

Although all the experiments are from a biology background, the scope of BCD is not confined to it. It can be applied to any longitudinal/spatial dataset affected by batch effects where the temporal/spatial locality holds. BCD also illustrates that batch the effect correction problem is related to the alternative clustering problems. Over the past few years, many advanced alternative clustering models have been introduced, and translate them to this context may result in better performance.

## 5 Summary

We defined a novel pairwise distance of the cells, namely Batch-Corrected Distance (BCD), where the effect of the unwanted clustering is controlled. Results show our method achieves more accurate clusters and better visualizations than state-of-the-art methods on longitudinal datasets. The BCD can be directly integrated into most clustering and visualization methods to enable more scientific findings.

## 6 Acknowledgements

This work was supported in part by grant number 2018-182735 to KC, Human Breast Cell Atlas Seed Network Grant (HCA3-0000000147) to KC from the Chan Zuckerberg Initiative DAF, an advised fund of Silicon Valley Community Foundation, grant RP180248 to KC from Cancer Prevention & Research Institute of Texas, The University of Texas MD Anderson Cancer Center Pre-Cancer Atlas Project to KC, and The University of Texas MD Anderson Cancer Center Colorectal Cancer Moonshot and P30 CA016672 (US National Institutes of Health/National Cancer Institute) to the University of Texas Anderson Cancer Center Core Support Grant.

## Notes

https://github.com/KChen-lab/varactor

### Competing Interest Statement

The authors have declared no competing interest.

